# Nonlinear relationship between multimodal adrenergic responses and local dendritic activity in primary sensory cortices

**DOI:** 10.1101/814657

**Authors:** Yair Deitcher, Yonatan Leibner, Sara Kutzkel, Neta Zylbermann, Michael London

## Abstract

The axonal projections of the adrenergic system to the neocortex, originating from the locus coeruleus (LC), form a dense network. These axons release the neuromodulator norepinephrine (NE) which is involved in many cognitive functions such as attention, arousal, and working memory. Using two-photon Ca^2+^ imaging of NE axons in the cortex of awake mice, we investigated what drives their phasic activity. We discovered that NE axons in the primary somatosensory cortex responded robustly and reliably to somatosensory stimulation. Surprisingly, the same axons also responded to stimuli of other modalities (auditory and visual). Similar responses to all three modalities were observed in the primary visual cortex as well. These results indicate that phasic responses of NE axons to sensory stimuli provide a robust multimodal signal. However, despite the robustness, we also noticed consistent variations in the data. For example, responses to whisker stimulations were larger than to auditory and visual stimulations in both the barrel and the visual cortices. To test whether the variations in NE axonal responses can carry behaviorally meaningful information, we trained mice in an associative auditory fear conditioning paradigm. We found that following conditioning the response of NE axons increased only for CS+, namely the signal undergoes experience-dependent plasticity and is specific to meaningful sounds. To test if variations in NE axonal responses can differentially affect the cortical microcircuit, we used dual-color two-photon Ca^2+^ imaging and studied the relationship between the activity of NE axons and local dendrites. We found dendritic Ca^2+^ signals in barrel cortex in response to auditory stimuli, but these responses were variable and unreliable. Strikingly, the probability of such dendritic signals increased nonlinearly with the Ca^2+^ signals of NE axons. Our results demonstrate that the phasic activity of the noradrenergic neurons may serve as a robust multimodal and plastic signal in sensory cortices. Furthermore, the variations in the NE axonal activity carry behaviorally meaningful signals and can predict the probability of local dendritic Ca^2+^ events.

## Introduction

Neuromodulation plays an important role in cortical processing and sensory integration. A dominant neuromodulator in the cerebral cortex is norepinephrine (NE) secreted by neurons of the locus coeruleus (LC), a small brain stem nucleus comprised of only ~1,500 neurons in rodents (Swanson 1976). LC neurons fire tonically and their firing rate correlate with brain states. LC neurons also show phasic evoked activity patterns in response to salient stimuli. These LC neurons are active in various conditions such as in sleep-wake transitions (Aston-Jones and Bloom 1981b; Carter et al. 2010), during specific aspects of learning (Sara and Segal 1991), and during novel sensory stimulation (Hervé-Minvielle and Sara 1995). LC neurons project widely all over the brain, including to the cerebral cortex. However, it is not clear if they form a homogeneous population which projects to all cortical regions or perhaps form subpopulations with specific projection patterns. Experimentally there are conflicting results. On the one hand, tracing methods found that the LC is a relatively homogeneous structure in which neurons receive similar inputs from many sources and send output broadly to the similar brain regions (Schwarz et al. 2015; Kim et al. 2016). On the other hand, other studies showed heterogenous organization of the LC demonstrating selective projection neurons to distinct cortical areas (Chandler et al. 2014; Waterhouse and Chandler 2016). Debate continues about the homogenous or heterogeneous nature of LC axons, and the specificity of their projections to the cerebral cortex, which may have significant functional consequences.

NE has been shown to modulate sensory processing in multiple ways, including selective gating, improvement of signal-to-noise ratio and sharpening of neurons tuning curves (Berridge and Waterhouse 2003; Manunta and Edeline 2004; Sara 2009; Polack et al. 2013). The main method for deducing NE activity is to record the spiking activity of LC neurons (Aston-Jones and Bloom 1981b). However, NE-release from axons in the cortex might not scale linearly, or even monotonically with the firing rate of LC neurons (Florin-Lechner et al. 1996). Furthermore, only a sub-population of LC neurons project to the neocortex (Schwarz et al. 2015; Kim et al. 2016; Rho et al. 2018), thus recording from LC neurons is not sufficient for understanding the activity of NE fibers in sensory cortices.

To detect in situ NE secretion, several studies used in vivo electrochemical measurements of NE. However, these methods still suffer from limitations such as the time scale of voltammetric detection and selectivity to NE (Park et al. 2018). Recent studies used two-photon Ca^2+^imaging to directly measure activity of NE axons in the cortex and to correlate their activity with pupil diameter and locomotion (Reimer et al. 2016; Breton-Provencher and Sur 2018). Others have developed optical imaging methods to measure the binding of NE to various fluorescent indicators (Muller et al. 2014; Dunn et al. 2018; Feng et al. 2019). However, little is known about the sensory cues that induce phasic NE-release in sensory cortex and the consequences of this release on synaptic integration in dendrites and spines, where the adrenergic receptors are found (Herkenham 1987; Nicholas et al. 1993; Aoki et al. 1998; Wang et al. 2007).

Here, we examined the conditions leading to evoked responses of NE axons in two sensory cortices. Using two-photon Ca^2+^imaging in awake mice, we found that NE axons in the barrel and visual cortices responded to stimuli of various modalities, including somatosensory, auditory and visual. We further show that the NE axonal responses to auditory stimulation in the barrel cortex exhibit associative plasticity, following acquisition of associative auditory fear conditioning. Finally, to examine directly the relationship between the axonal activity of NE projections and cortical dendritic activity, we performed dual-color two-photon Ca^2+^ imaging of local dendrites and NE axons. Interestingly, we found that local dendrites in the barrel cortex responded to auditory stimulation, and the probability of these events increases monotonically with the activity levels of NE axons. Our results are consistent with the hypothesis that NE can tune the excitability and sensitivity of dendrites to long range input projections (Phillips et al. 2016, 2018; Labarrera et al. 2018).

## Results

### Response of Ca^2+^ activity in LC axons to whisker stimulation in the barrel cortex

To examine the distribution of norepinephrine (NE) axons in the barrel cortex, we used selective immunostaining for the noradrenaline transporter (NET), which is expressed exclusively in noradrenergic axons (Lorang et al. 1994). We found dense axonal innervation throughout the neocortical layers in the barrel cortex (Figure 1A). To image the activity of adrenergic axons in the barrel cortex, we selectively expressed the genetically encoded Ca^2+^ indicator GCaMP6s (Chen et al. 2013) in the Locus Coeruleus (LC) of knock-in mice expressing Cre in tyrosine hydroxylase positive neurons (Lindeberg et al. 2004, Figure 1B). We validated the specificity of viral expression by comparing fluorescence in the LC and in the barrel cortex, with that of tyrosine hydroxylase (TH) and NET immunofluorescence, respectively (Figure 1C). Indeed, all axons expressing GCaMP6s were labeled with the NET antibody (1,770 regions of interest (ROIs), 3 mice).

**Figure 1.**
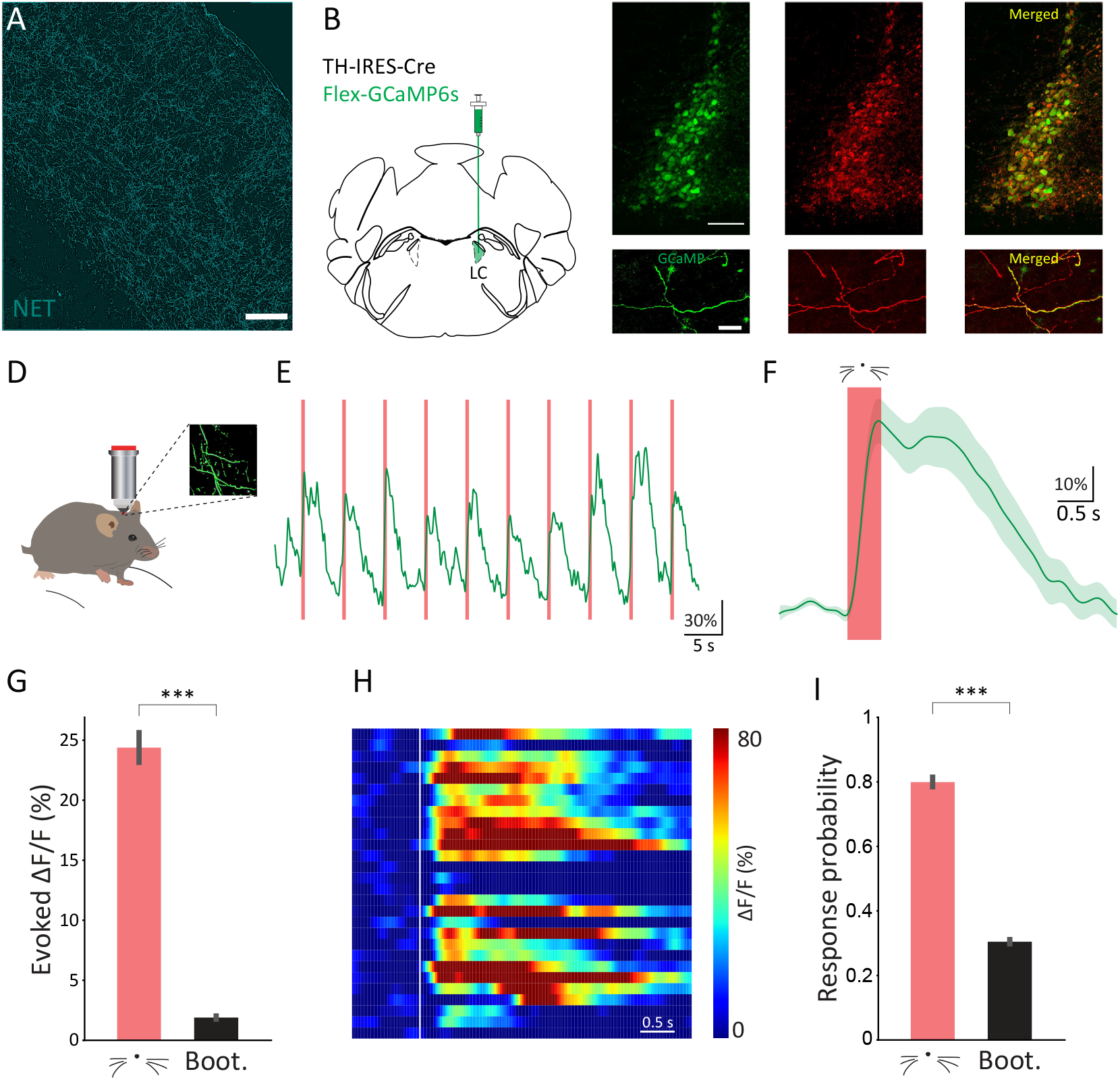
Ca^2+^ imaging of Locus Coeruleus axonal activity in the barrel cortex. **A.** Noradrenergic (NE) axons in the barrel cortex labeled using a noradrenaline transporter (NET) antibody (red). Scale bar, 200 μm. **B.** Stereotactic injections of Flex-GCaMP6s into the LC of TH-IRES-Cre mice. **C** Top: viral expression of GCaMP6s in the LC (green), tyrosine hydroxylase (TH) antibody staining (red), and a merged image (yellow). Scale bar, 100 μm. Bottom: NE axons in the barrel cortex expressing GCaMP6s (green), NET antibody staining (red) and a merged image (yellow). Scale bar, 20 μm. **D.** Schematic illustration of two-photon Ca^2+^ imaging configuration from head-restrained mice, walking on a wheel. Inset: average image showing LC axons expressing GCaMP6s. **E.** An example fluorescence trace of an LC axon in response to whisker stimulation. Red lines indicate the period of whisker stimulation. **F.** Mean Ca^2+^ response of the same LC axon to whisker stimulation (S.E.M in green shaded area) **G.** Comparison of the mean evoked response to whisker stimulation and the bootstrap results for all LC axons. **H.** Color raster plot of Ca^2+^ signals from all trials from the same axon, each row represents one trial. White line indicates the onset of the whisker stimulation. **I.** Comparison of response probability to whisker stimulation and bootstrap results for all LC axons.

We then imaged the adrenergic fibers in the barrel cortex in head-fixed awake mice (Figure 1D). In agreement with previous reports (Reimer et al. 2016; Breton-Provencher and Sur 2018), we found a correlation between the activity of NE axons and pupil diameter (Figure S1A-B, n = 16 ROIs, 4 Fields of View (FoV), 3 mice).

As sensory evoked response in NE axons have never been explored, we tested whether in the barrel cortex the NE axons would respond to air puff stimulation applied to the whiskers. We found evoked responses to whisker stimulation with a prominent increase in Ca^2+^ (Figure 1E-G, mean evoked response (for definition see Methods): 24.39% ± 1.25%, bootstrap: 1.9 ± 0.13%; n = 78 ROIs, 1,777 trials, 15 FoV, 4 mice; p < 0.001, t test). We validated that these responses were reflecting real signals and were not due to movements of the imaging plane by comparing the axonal responses to the change in signal over time of auto-fluorescent ‘blebs’ (Reimer et al. 2016, Figure S1C, axonal evoked response, 24.39% ± 1.25%, blebs evoked response 0.92 ± 0.33%, n = 78 axonal ROIs, n = 59 blebs ROIs, 15 FoV, 4 mice; p < 0.001, t test).

The axonal response appeared in single trials of stimulation (Figure 1H), with a response probability of 80%, indicating a reliable axonal response (Figure 1I, response probability: 0.8 ± 0.02; bootstrap: 0.3 ± 0.01; n = 78 ROIs, 1,777 trials, 15 FoV, 4 mice; p < 0.001, t test. Figure S1D, response probability axons: 0.8 ± 0.02; blebs: 0.09 ± 0.01; n = 78 axonal ROIs, n = 59 blebs ROIs, 15 FoV, 4 mice; p < 0.001, t test). We conclude that NE axons in barrel cortex display a robust and reliable response to whisker stimulations which can be detected even in individual trials.

### Response of LC axons to multiple stimuli in the barrel and visual cortices

Primary sensory areas are notoriously known for their neurons highly selective responses (Kerr et al. 2007; Niell and Stryker 2008). In contrast, anatomically it has been shown that LC neurons receive inputs from many sources (Schwarz et al. 2015), and indeed electrophysiological recordings from the LC found neurons which respond to stimuli of two or more sensory modalities (Aston-Jones and Bloom 1981a). We have found that LC axons in barrel cortex reliably respond to somatosensory stimulation. The question is whether axons of LC neurons projecting to primary sensory areas show selectivity to that modality or do they respond to stimuli of other modalities as suggested from the multimodal properties of LC neurons.

We therefore tested the responses to sensory stimuli from other modalities of LC axons in barrel cortex. We found that the same NE axons also responded to simple auditory and visual stimuli (see Methods) with a similar increase in Ca^2+^ signal (Figure 2A-B, Figure S2A-B). Given the recordings were made in a primary sensory area we were surprised by the robustness and reliability of these cross-modal responses. These results suggest that NE may act as a multimodal signal, conveying information to primary sensory cortices about other modalities.

**Figure 2.**
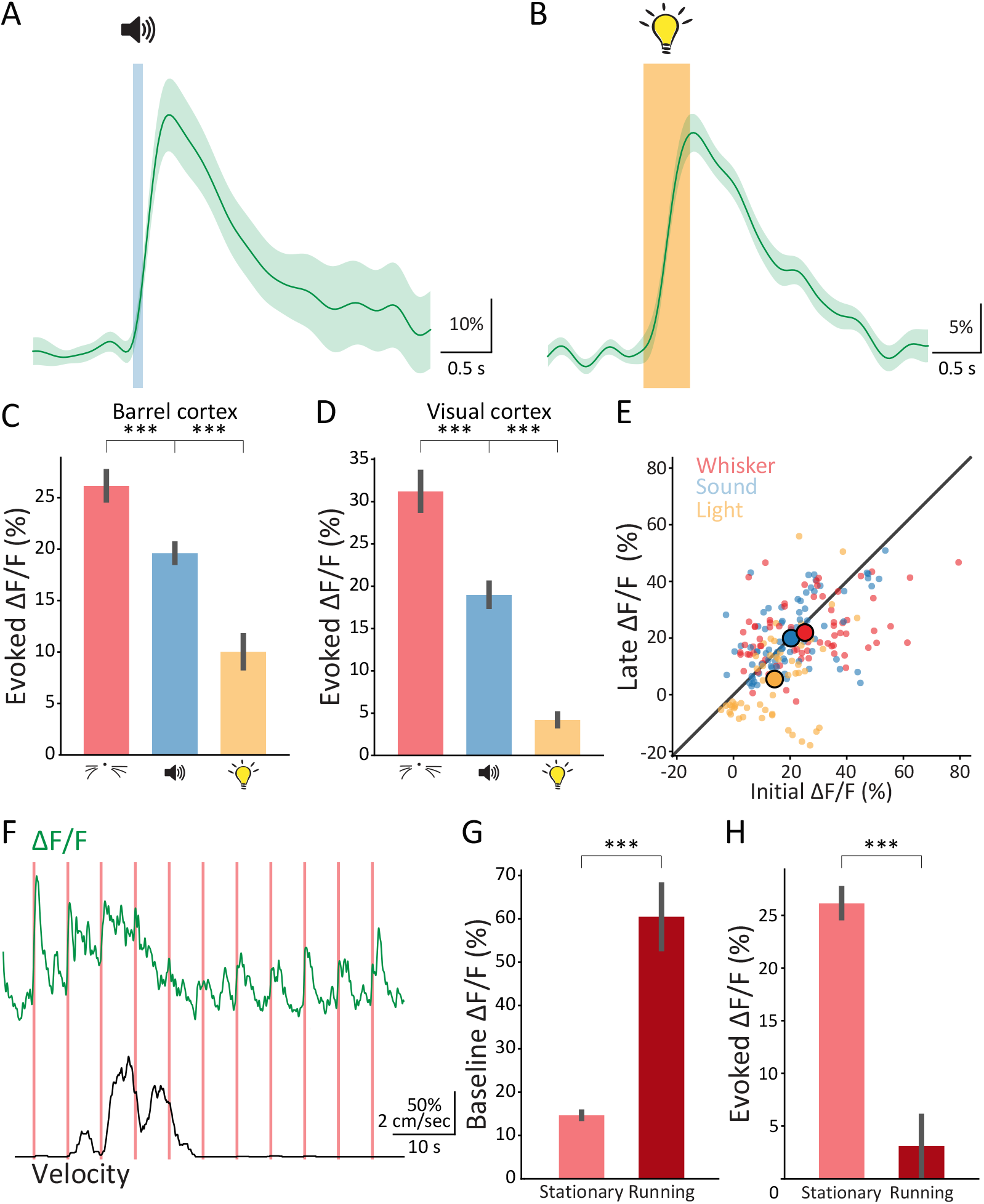
Response of LC axons to multiple stimuli modalities in the barrel and visual cortices. **A.** Mean Ca^2+^ response (solid line) of the same LC axon from Figure 1 (barrel cortex) to auditory stimulation. Blue shaded area indicates the time and duration of the auditory stimulation (SEM in shaded green) **B.** Mean Ca^2+^ response of the same axon to visual stimulation. Yellow shaded area indicates the time and duration of the visual stimulation. **C.** Comparison of the mean evoked response between the three conditions (whisker, auditory and visual stimulations) in the barrel cortex **D.** Comparison of the mean evoked response between the three conditions (whisker, auditory and visual stimulations) in the visual cortex. **E.** Comparison of the mean response of the five initial responses of LC axons and the mean response of the five latest responses. **F.** Top, Ca^2+^ fluorescence traces of an LC axon in response to whisker stimulation (green). Bottom, velocity of the mouse (black). Red shaded areas indicate whisker stimulations. **G.** Comparison of the baseline activity in LC axons in the barrel cortex during stationary and running epochs. **H.** Comparison of the mean evoked response in LC axons in the barrel cortex during stationary and running epochs.

However, despite the robustness, we also noticed variations in the responses. When comparing the response of LC axons in barrel cortex to the different stimuli, we found that the largest response was to the whisker stimulation, then to the auditory stimulation, and the weakest response was induced by the visual stimulation (Figure 2C, whisker: 26.16% ± 1.38%, n = 78 ROIs, 1,777 trials, 15 FoV, 4 mice; auditory: 19.61% ± 0.89%, n = 92 ROIs, 2,707 trials, 15 FoV, 4 mice; visual: 10.02% ± 1.57%, n = 53 ROIs, 1,031 trials, 8 FoV, 3 mice; p < 0.001; one-way ANOVA, Tukey’s multiple comparison test). Similar results were found when comparing the response probability of the different conditions (Figure S2C, whisker: 0.8 ± 0.01, n = 78 ROIs, 1,777 trials, 15 FoV, 4 mice; auditory: 0.71 ± 0.02, n = 92 ROIs, 2,707 trials, 15 FoV, 4 mice; visual: 0.42 ± 0.04, n = 53 ROIs, 1,031 trials, 8 FoV, 3 mice; p < 0.001, one-way ANOVA, Tukey’s multiple comparison test).

Is this multimodal response specific to the barrel cortex or is this phenomenon similar across sensory cortices? To directly address this question, we imaged NE axons in the visual cortex and presented the mice with an identical set of sensory stimuli. Similar to the barrel cortex, in the visual cortex as well we found multimodal responses to whisker, auditory, and visual stimuli (Figure 2D, whisker: 31.22% ± 2.27%, n = 34 ROIs, 573 trials, 7 FoV, 2 mice; auditory: 18.99% ± 1.38%, n = 44 ROIs, 709 trials, 10 FoV, 2 mice; visual: 4.21% ± 0.7%, n = 24 ROIs, 367 trials, 6 FoV, 2 mice; p < 0.001, one-way ANOVA, Tukey’s multiple comparison test). Similar results were also found when comparing the response probability of the different conditions (Figure S2D, whisker: 0.83 ± 0.02, n = 34 ROIs, 573 trials, 7 FoV, 2 mice; auditory: 0.64 ± 0.03, n = 44 ROIs, 709 trials, 10 FoV, 2 mice; visual: 0.33 ± 0.04, n = 24 ROIs, 367 trials, 6 FoV, 2 mice; p < 0.001, one-way ANOVA, Tukey’s multiple comparison test).

Remarkably, the LC axons in visual cortex responded in a similar manner to those in the barrel cortex, namely, the weakest response was to the visual stimulation, a larger response to the auditory stimulation and the largest response was to whisker stimulation. These results may suggest that the variations in the axonal responses reflects the salience of the stimulation, independent of the modality of the primary sensory cortex where the axons project.

To test if the response of LC axons to a series of stimuli shows interesting dynamics (adaptation or facilitation), we compared the initial responses (the mean response to the first five trials with sensory stimulus) to the late responses (the mean response to the last five trials). Almost no change appeared in all three modalities (Figure 2E). As startle responses are highly adaptive (Ding et al. 2013) our results indicate that the NE axonal activity most likely does not represent a startle response.

As shown above, we found variations in the NE axonal responses to stimuli from different modalities. We next examined if there are variations also in response to the same stimulus, depending on the behavioral state, namely, when the animal is running or standing. We therefore analyzed the response of LC axons in the barrel cortex to whisker stimulation while the mouse was running or standing. Interestingly, when whisker stimulation was presented during epochs where the mouse was running, there was a very small axonal response compared to the response when the stimulus was presented while the mouse was stationary (Figure 2F-H, whisker_stat_: 26.16% ± 1.38%, n = 78 ROIs, 1,777 trials, 15 FoV, 4 mice; whisker_run_: 3.1% ± 2.88%, n = 20 ROIs, 32 trials, 5 FoV, 3 mice; p < 0.001, t test). Similar results were found for auditory and visual stimulations (auditory_stat_: 19.61% ± 0.89%, n = 92 ROIs, 2,707 trials, 15 FoV, 4 mice, auditory_run_: −6.07% ± 1.63%, n = 26 ROIs, 65 trials, 6 FoV, 3 mice, p < 0.001, t test; visual_stat_: 10.02% ± 1.57%, n = 53 ROIs, 1,031 trials, 8 FoV, 3 mice, visual_run_: 3.53% ± 1.76%, n = 35 ROIs, 69 trials, 5 FoV, 2 mice; p < 0.01, t test). As previously reported (Reimer et al. 2016), at the beginning of walking the activity of NE axons increases resulting in a relative large shift in baseline activity (Figure 2G). Thus, the decrease in response during running (Figure 2H) may result from a saturation in the Ca^2+^ sensor itself (ceiling effect) or alternatively reflect a more general phenomenon where sensory stimulation during active states results in weaker responses (Petersen et al. 2003).

### Experience dependent plasticity in NE axons activity following associative fear

Our data demonstrate that the LC axonal activity in sensory cortices conveys a robust multimodal spatially-diverse signal. The data presented in figure 2 also hint that some information is carried at a finer scale of variation. First, there are consistent variations in the responses to stimuli of different modalities (Figure 2C, D). Second, the responses to the same stimulus change between behavioral states (Figure 2F, H). Are the responses of LC axons capable of showing experience dependent plasticity such that they can deliver information on sensory cues at this finer resolution (beyond the mere appearance of a salient sensory stimulus)?

To directly address this question, we trained mice in an associative auditory fear conditioning (FC) paradigm and measured the freezing behavior of mice in response to the conditioned stimulus (CS+) as compared to a non-associated stimulus (CS–; not paired with a foot shock, Figure 3A). Notably, under freely behaving conditions, the mice showed a prominent and discriminative fear response to the CS+ (Figure 3B, CS^−^ freezing: 20.86% ± 2.97%, CS^+^_freezing_: 71.75 ± 5.11%, n = 21 mice; p < 0.001, paired t test), indicating successful fear memory learning. To estimate fear expression in head-fixed mice, we measured the change in pupil diameter in response to the auditory cues (CS^−^ versus CS^+^) during habituation and recall (Abs et al. 2018; Garcia-Junco-Clemente et al. 2019, Figure 3C). Following fear conditioning, the pupil evoked responses increased for the CS^+^ but not for the CS^−^ (Figure 3D-F; CS^−^_before_: 0.46% ± 1.46%, CS^+^_before_: −0.57% ± 0.62%; n = 7 mice, 263 trials; p = 0.47, paired t test; CS^−^_after_: 1.88% ± 0.45%, CS^+^_after_: 10.2% ± 2.15%, n = 9 mice, 352 trials; p < 0.01 paired t test). Thus, pupil response can be used as a faithful readout of fear conditioning.

**Figure 3.**
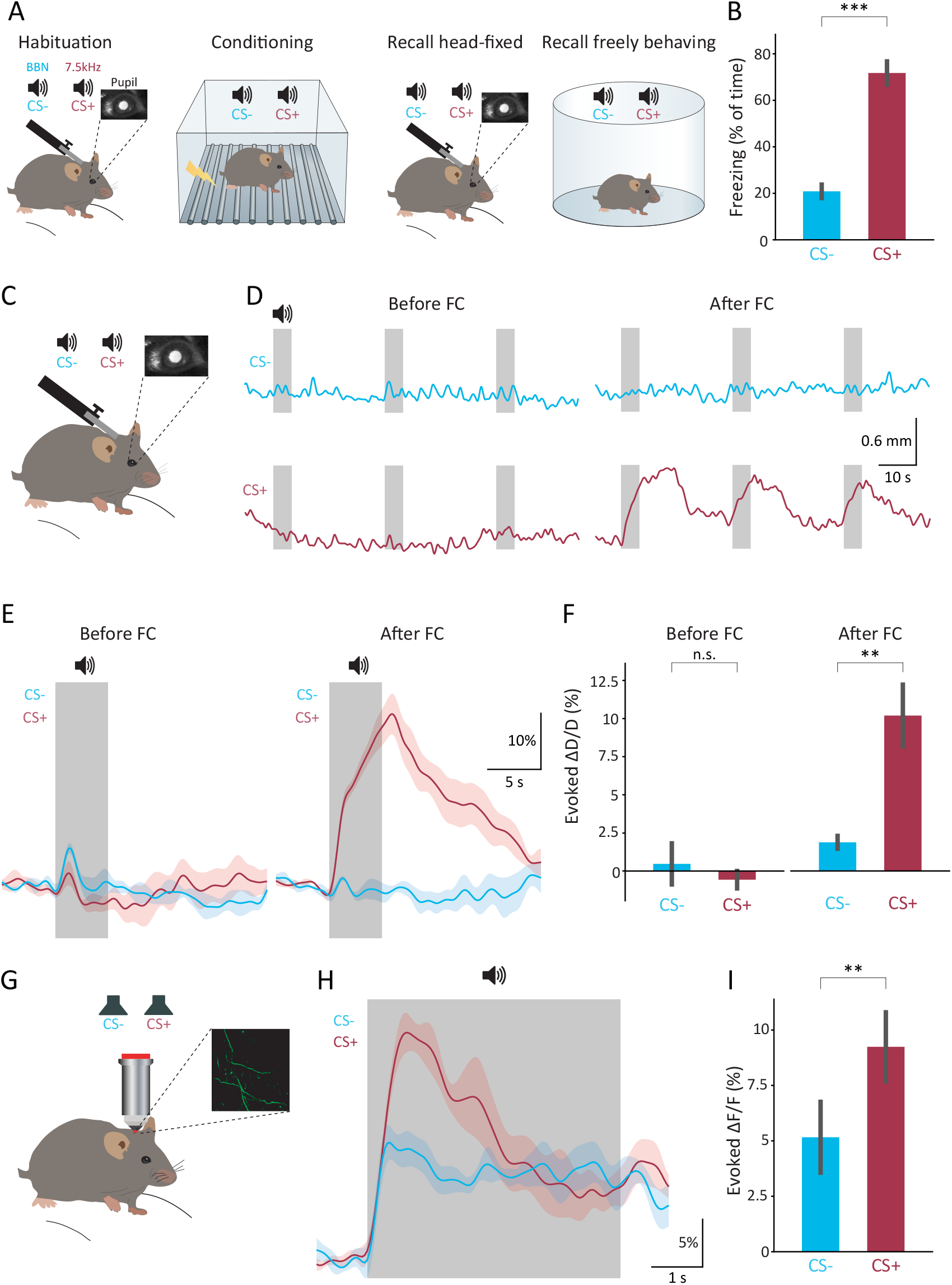
Experience dependent plasticity in LC-axon activity following associative fear acquisition. **A.** Schematic illustration of the associative auditory fear conditioning paradigm. **B.** Comparison of the freezing behavior (in a freely behaving memory recall session) to the conditioned stimulus (CS+) and to a non-conditioned stimulus (CS–; not paired with a foot shock). **C.** Schematic illustration of a memory recall session in a head-fixed mouse. **D.** Examples of pupil diameter responses to CS^−^ (blue) and CS^+^ (red) before (left) and after (right) fear conditioning (FC). Gray shaded area indicates the CSs presentations. **E.** Mean pupil response to the CSs before (left) and after (right) FC in one mouse. Red- and blue-shaded areas are the S.E.M. **F.** Comparison of the mean evoked pupil response before (left) and after (right) FC in all mice. **G.** Schematic illustration of two-photon Ca^2+^ imaging in head-restrained mice during memory recall. **H.** Mean Ca^2+^ response of LC axons to CSs during memory recall in one mouse. **I.** Comparison of the mean evoked Ca^2+^ response to the CSs in all mice.

To test whether NE axons in the barrel cortex show a discriminative response following auditory fear conditioning we combined the fear conditioning behavior with in vivo two-photon Ca^2+^ imaging of LC axons (Figure 3G). During fear memory recall, NE axons showed a larger Ca^2+^ increase for the CS^+^ than for the CS^−^ (Figure 3H-I, CS^−^: 5.16% ± 1.64%, CS^+^: 9.24% ± 1.6%, n = 18 ROIs, 168 trials, 11 FoV, 3 mice; p < 0.01 paired t test). Consistent with this result, mice that experienced fear conditioning but failed to form a stable memory, showed no significant difference in Ca^2+^ response following both CSs (Figure S3A, CS^−^: 2.31% ± 0.92%, CS^+^: 2.61% ± 1.04%, n = 22 ROIs, 211 trials, 8 FoV, 2 mice; p = 0.72 paired t test), further supporting the interpretation that the potentiation in LC axonal responses resulted from the association of the CS+ auditory stimulus with fear. These data demonstrate that recall of an aversive auditory memory is associated with a selective pronounced Ca^2+^ increase in NE axons located in the cortex. More generally, these results show that the robust multimodal spatially-diverse signal of the cortical projections of the NE system, is plastic and can modify the information it conveys in a behaviorally meaningful manner.

### Nonlinear relationship between auditory evoked LC axonal and local dendritic responses in barrel cortex

Having established evoked responses of NE axons activity in the barrel cortex, we next aimed to explore the relationship between noradrenergic axon transient responses and the local circuit activity. Previously it has been suggested that the excitability of dendrites located in the upper layers of the cortex is subject to neuromodulation (Barth et al. 2008; Dembrow et al. 2010; Labarrera et al. 2018; Williams and Fletcher 2019). We thus expressed the red-shifted genetically encoded calcium indicator, jRGECO1a (Dana et al. 2016), in neurons located in the barrel cortex and GCaMP6s in the LC (Figure 4A-B). This allowed us to perform dual-color functional imaging, of both LC axons and dendritic branches of neurons in layer 1 of the barrel cortex. Imaging of spontaneous activity showed that Ca^2+^ events in NE axons were correlated with Ca^2+^ events in the local dendrites (Figure 4C). The cross-correlation function between LC axons activity and the mean dendritic activity, showed a large peak close to zero lag (Figure 4D, 15 axonal ROIs, 52 dendritic ROIs, 5 FoV, 3 mice). While this large peak at this lag indicates a strong temporal correlation between the two signals, due to the different rise and decay time-constants of jRGECO1a (Dana 2016) and GCaMP6s (Chen et al. 2013), we were unable to determine the precise temporal relationship between the NE axonal and local dendritic activity.

**Figure 4.**
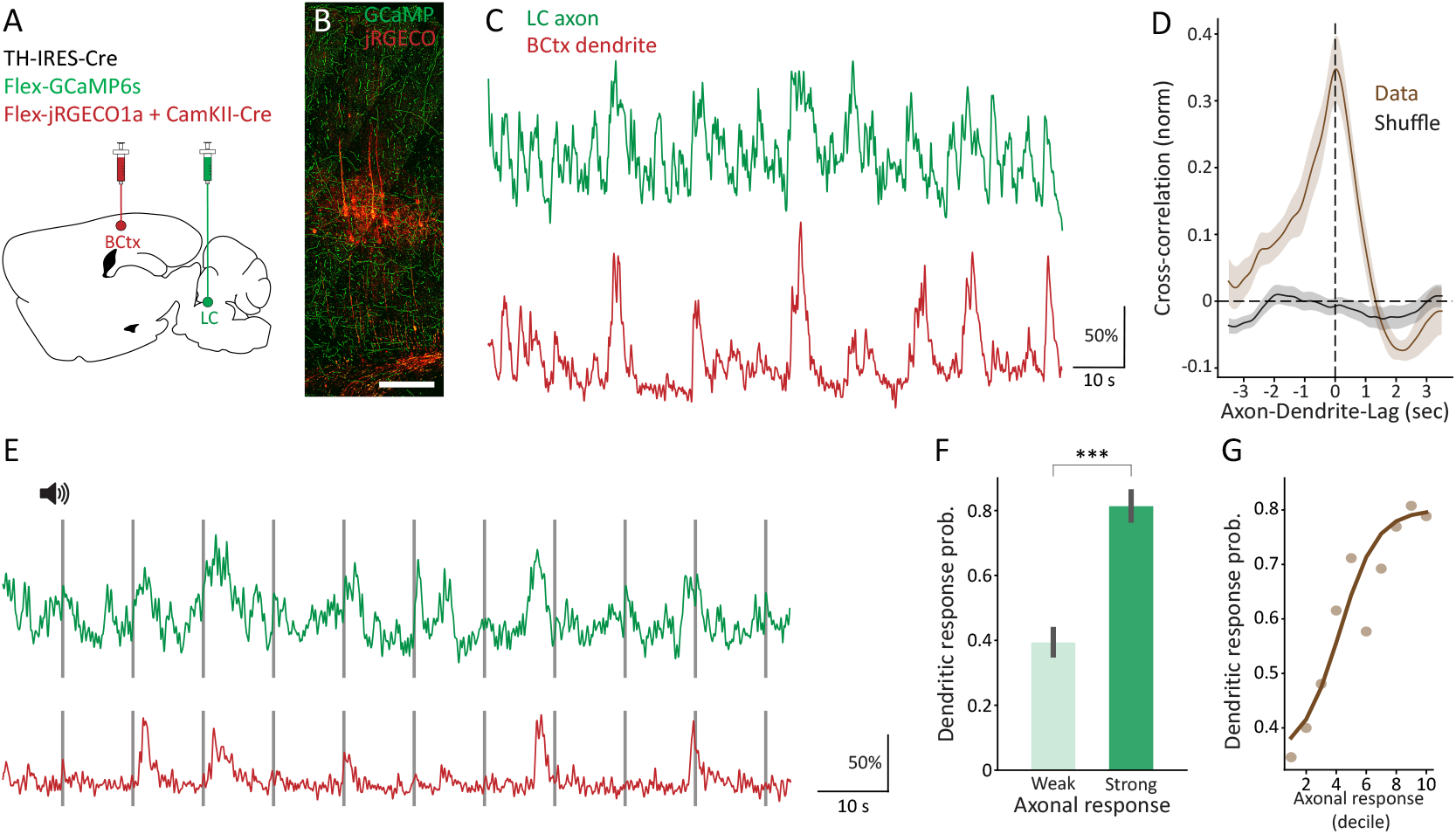
Relationship between the sensory responses of LC axons and local barrel cortex dendrites. **A.** Stereotactic injections of Flex-jRGECO1a mixed with CamKII-Cre into the barrel cortex and Flex-GCaMP6s into the LC of TH-IRES-Cre mice. **B.** Noradrenergic axons in the barrel cortex expressing GCaMP6s (green) and neurons in the barrel cortex expressing jRGECO1a (red). Scale bar, 200 μm. **C.** Dual-color two-photon imaging of NE axons and local dendrites. Example Ca^2+^ fluorescence traces of an LC axon (green) and of a local dendrite (red). **D.** Mean cross-correlation between the mean activity of LC axons and the mean activity of an adjacent dendrites (brown). Same analysis for shuffled data in black. Shaded areas are the S.E.M. **E.** Example Ca^2+^ fluorescence traces of an LC axon (green) and of a local dendrite (red) in response to auditory stimulation. Gray shaded area indicates the period of the auditory stimulation. **F.** Comparison of the mean dendritic response probability when axonal response is weak and strong. **G.** The mean dendritic response probability as a function of the axonal response. Dark brown line is a sigmoid fit.

As we have shown above, LC axons in the barrel cortex also respond to auditory stimulation (Figure 2). Thus, we tested the response of local dendritic branches to auditory stimulation and found that a subset of these dendrites indeed responded to auditory stimuli as well. However, the response reliability of these dendrites was not very high and they did not respond in all trials of stimulation (Figure 4Es). To check if variations in NE activity could explain some of this unreliability, we compared the response probability of a dendrite when the axonal response was weak (1^st^ quartile; see Methods) and when the axonal response was strong (4^th^ quartile). Notably, the dendritic response probability was larger when the axonal response was strong (Figure 4F, weak response: 0.39 ± 0.04, strong response: 0.81 ± 0.04, n = 26 dendritic ROIs, 27 axonal ROIs, 356 trials, 9 FoV, 3 mice; p < 0.001, paired t test; Figure S4A). Moreover, the mean evoked dendritic response was also larger when axonal response was large (Figure S4B, weak response, 4.54% ± 0.8%, strong response, 11.56% ± 1.28%, n = 26 dendritic ROIs, 27 axonal ROIs, 356 trials, 9 FoV, 3 mice; p < 0.001, paired t test; See also Figure S4C for quantification of the wide dynamic range of the axonal fluorescent signal).

Finally, we examined the response probability of these barrel cortex dendrites as a function of the NE axonal response (Figure 4G). Strikingly, as the axonal response increased so did the probability of the dendrites to respond (to auditory stimuli). This relationship is not linear, showing saturation at both low and high ends and was fitted with a sigmoidal function. These data indicate that dendritic response probability in the barrel cortex to an auditory stimulation, increases when NE activity is higher, suggesting that the transient variations in NE concentration in the cortex following a sensory stimulation may explain part of the dendritic responses.

## Discussion

In this study, we examined the activity of noradrenergic LC axons in primary sensory cortices. We imaged the Ca^2+^ activity of NE axons in awake mice and found a multimodal response to somatosensory, auditory and visual stimulations in the barrel cortex. Similar results were observed also in the visual cortex. These results suggest that NE acts as a multimodal spatially diverse signal, conveying integrated information from distinct sensory modalities to different areas of the sensory cortex. Moreover, the auditory response can be modulated following an associative auditory fear conditioning experience. Namely, NE axons show a selective Ca^2+^ increase for a conditioned stimulus (CS^+^) after it was paired with a foot shock. Finally, we simultaneously investigated the responses of LC axons and local dendrites in the barrel cortex to auditory stimulation and found that the response probability of barrel cortex dendrites to auditory stimulation, increases monotonically and nonlinearly with the Ca^2+^ activity of noradrenergic LC axons.

### NE axons show multimodal spatially-diverse reliable response

The noradrenergic system is multimodal in various respects. Electrophysiological recordings in the LC of rats have shown responses to simple stimuli of different modalities (Aston-Jones and Bloom 1981a). In addition, anatomical studies (Kim et al. 2016) have shown that individual LC neurons may project to more than one primary sensory area. Thus, LC neurons receive input from and respond to various modalities and project to various targets where sensory information is processed. Our results are consistent with these findings, and show functionally that LC axons in primary sensory cortices respond to sensory stimuli of various modalities. This may suggest that NE activity is a nonspecific broad signal that conveys little information. However, the details of the responses of NE signals as we observe, show interesting variations to behavioral states (Figure 2H), to the modality or saliency of the stimulus (Figure 2C) and also plasticity that corresponds to behavioral importance (Figure 3I). Moreover, the variation in the NE signal predicts in a nonobvious way the activity of neurons in the local circuit (Figure 4G). Therefore, these data suggest that understanding the contribution of the NE signals to cortical processing requires high resolution sensitivity to small nuances in the NE signal variations.

### Relationship between spontaneous NE activity and cortical circuit activity

Most of what is known about the relationship between NE and activity of local cortical circuits relies on using pupil diameter measurements as a proxy to NE activity (Aston-Jones and Cohen 2005; Varazzani et al. 2015; Joshi et al. 2016; Reimer et al. 2016). Simultaneous recordings of neuronal activity in auditory cortex with pupil diameter, shows that brief pupil microdilations were associated with 5–20 mV depolarization in the membrane potential of cortical neuronal (McGinley et al. 2015). In both visual and somatosensory cortices, small spontaneous fluctuations in pupil diameter track changes in the intracellular dynamics of L2/3 neurons (Reimer et al. 2014). Our results on the high correlation between spontaneous NE axons activity and local dendritic activity is consistent with these results.

### NE signals and dendritic computation in perceptual sensitivity

NE is crucial for maintaining cognitive brain functions such as perception, attention, and learning (Berridge and Waterhouse 2003; Aston-Jones and Cohen 2005; Sara 2009; Sara and Bouret 2012). A recent study demonstrated that optogenetic stimulation of LC neurons, improves perceptual sensitivity in rats (Rodenkirch et al. 2019) and other reports showed that LC activation accelerates perceptual learning in the auditory system (Martins and Froemke 2015; Glennon et al. 2019). Moreover, there is evidence for noradrenergic involvement in perceptual shifts in human subjects (Einhäuser et al. 2008). However, there is little mechanistic understanding of how NE changes contribute to these effects on perception at the circuit and cellular level. One candidate mechanism is the contribution of NE to synergistic integration of information from different sources through its effect on dendritic excitability (Barth et al. 2008; Phillips et al. 2016, 2018; Labarrera et al. 2018).

Several studies in awake mice indicate that Ca^2+^ activity in the dendrites of cortical pyramidal cells is elevated during cognitive tasks, and that manipulating the activity of these dendrites shifted the perceptual threshold (Xu et al. 2012; Cichon and Gan 2015; Manita et al. 2015; Miyamoto et al. 2016; Takahashi et al. 2016; Ranganathan et al. 2018). Here, we show that the probability of a dendritic response to auditory stimulation increases monotonically with the activity levels of NE axons, suggesting that NE can act as a signal that lowers the threshold for the generation of dendritic Ca^2+^ events. Our study is in line with the proposal that large Ca^2+^ events in the dendrites are tuned by NE and that the sensitivity of the dendrites is modulated by NE (Phillips et al. 2016, 2018; Labarrera et al. 2018). Future experiments aimed at establishing this hypothesis will ideally include recordings from both NE signals and local dendritic activity in awake, behaving animals while they engage in a perceptual task.

## Supplementary figures

**Supplementary Figure 1.**
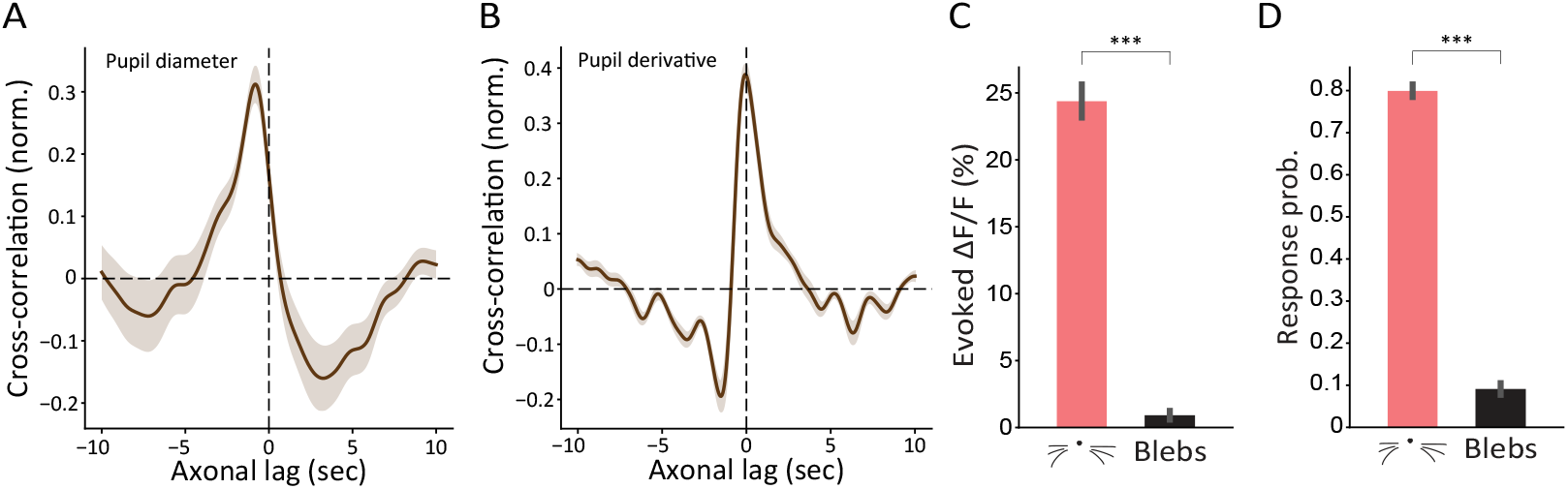
Correlation between pupil diameter and NE axons. Comparison of NE axons response and blebs response to whisker stimulation in the barrel cortex. **A.** Cross-correlation between pupil diameter and NE axons. **B.** Cross-correlation between pupil diameter derivative and NE axons. **C.** Comparison of the mean evoked response of NE axons and blebs to whisker stimulation. **D.** Comparison of the response probability of NE axons and blebs to whisker stimulation.

**Supplementary Figure 2.**
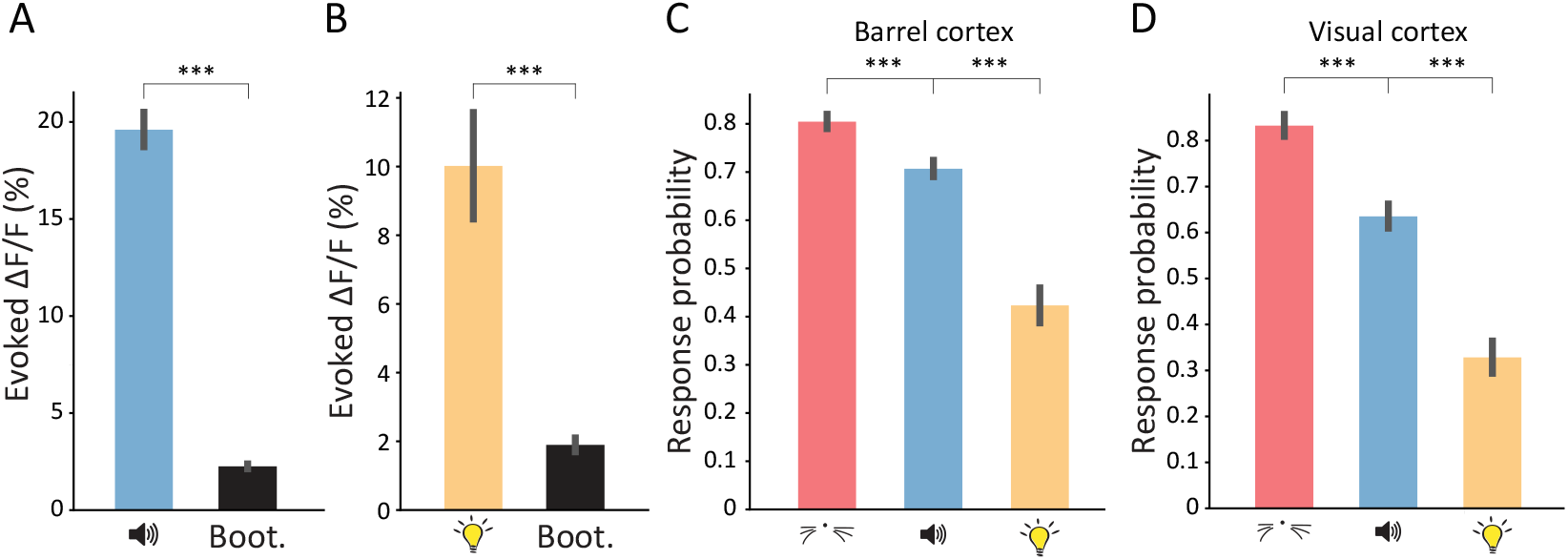
Response of LC axons to multiple stimuli in the barrel and visual cortex. **A.** Comparison of the mean evoked response to auditory stimulation and the bootstrap results for all LC axons. **B.** Comparison of the mean evoked response to visual stimulation and the bootstrap results for all LC axons. **C.** Comparison of the response probability between the three conditions (whisker, auditory and visual stimulations) in the barrel cortex. **D**. Same as C but in the visual cortex.

**Supplementary Figure 3.**
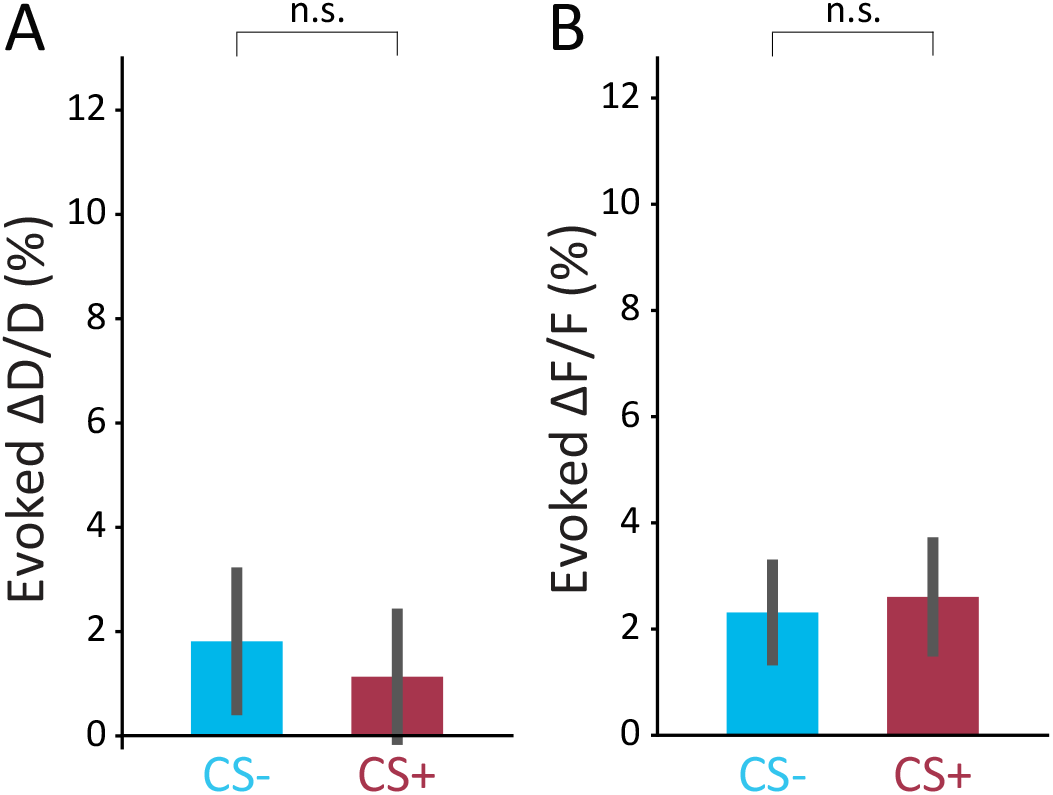
Pupil and NE axonal responses to auditory cue in mice that experienced fear conditioning but failed to form a stable memory. **A** Comparison of the mean evoked pupil response in mice that failed to form a stable memory. **B.** Comparison of the mean evoked Ca^2+^ response in mice that failed to form a stable memory.

**Supplementary Figure 4.**
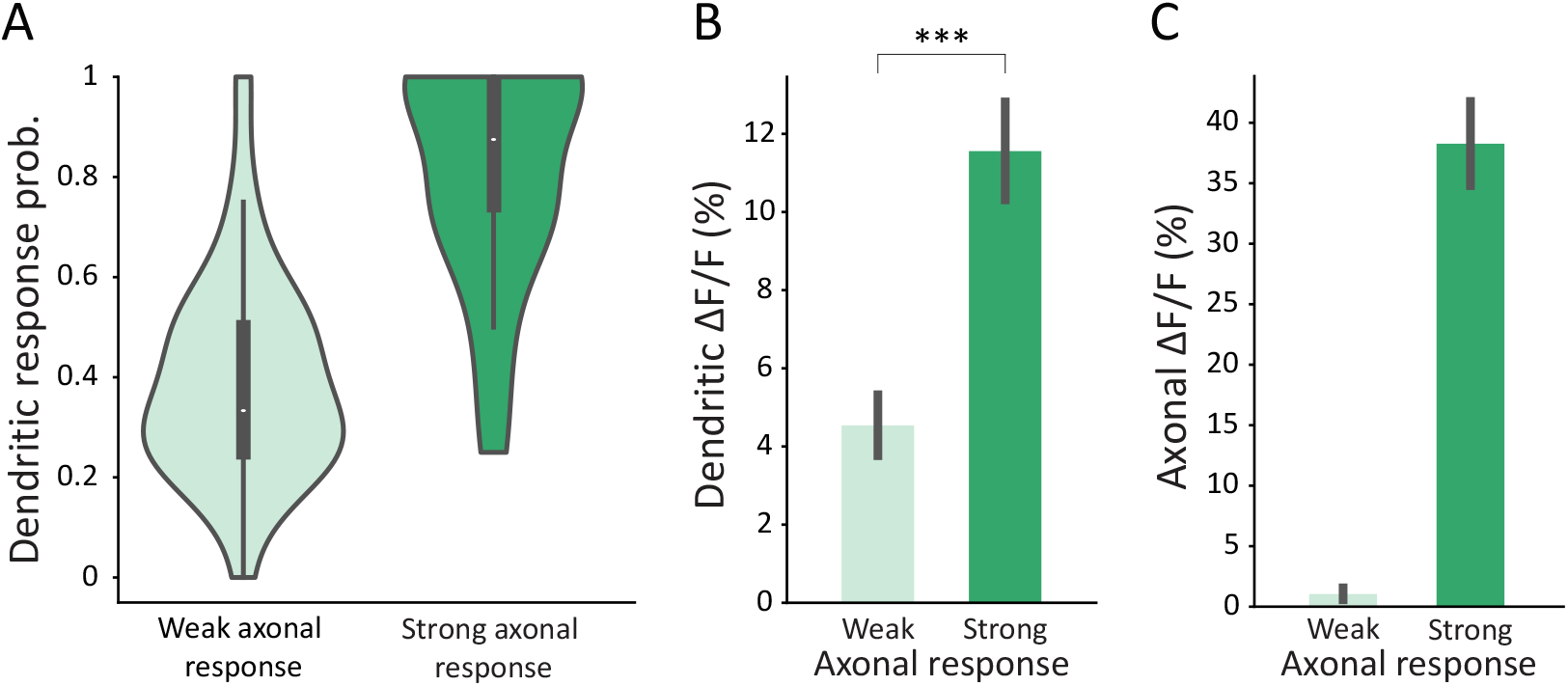
Comparison of dendritic response probability during weak and strong axonal responses. **A.** Violin plot showing the distribution of dendritic response probability when axonal response is weak or strong. The small black box shows the quartiles of the dataset and the whiskers extend to show the rest of the distribution. **B.** Comparison of the mean dendritic evoked response when axonal response is weak and strong. **C.** Comparison of the mean axonal evoked response when axonal response is weak (1.06% ± 0.5%) or strong (38.28% ± 3.55%).

## Acknowledgments

Inbal, Michael (pupil software), Tali kimchi

## Methods

### Animals

We used TH-IRES-Cre mice (Lindeberg et al. 2004), 8–13 weeks old. The Hebrew University Animal Care and Use Committee approved all experiments.

### Surgery and Viral Vector Injections

Viral vectors were delivered using standard stereotactic injections to the right locus coeruleus (Carter 2010, anteroposterior: −5.45 mm; mediolateral: 1.28 mm; dorsoventral: 3.65 mm) under isoflurane anesthesia. Expression of GCaMP6s (Chen et al., 2013) in LC neurons was achieved via viral injection of pAAV.CAG.Flex.GCaMP6s.WPRE.SV40 (AAV9) (Addgene). For sparse labeling of neurons in the barrel cortex with jRGECO1a (Dana 2016), pAAV.CAG.Flex.NES-jRGECO1a.WPRE.SV40 (AAV1; Addgene) was co-injected with diluted (1:10) AAV9.CamKII.Cre. A head-post was installed after viral injection.

Four weeks after the virus injection, a 3-mm craniotomy was drilled and a cranial window consisting of two stacked 3-mm coverslips under a 5-mm coverslip were sealed into the craniotomy.

### Two-Photon Calcium Imaging

Imaging from awake animals was performed with a low-power temporal oversampling (LOTOS) two-photon microscope (LotosScan2015, Suzhou Institute of Biomedical Engineering and Technology; http://english.sibet.cas.cn/) at 920 nm with a Ti:Sapphire laser (Vision II, Coherent, CA) and imaged through a 25x, 1.05 numerical aperture (NA) water immersion objective (Olympus, Japan). Frame images were acquired at 20-40 frames per second.

For dual-color imaging (Figure 4) a single excitation source at 1000 nm was used to excite both indicators.

### Fear conditioning

After mice were habituated to head-fixation under the microscope, both CSs (CS+: pure tone, 7.5 kHz; CS−: broadband noise (BBN)) were presented for habituation of the CSs. Each CS was presented 18-24 times (each presentation 5 sec long, 25 sec inter-trial interval). CS+ and CS− were presented in an alternating fashion.

The Fear Conditioning (FC) apparatus consisted of a conditioning box (18×18×30 cm), with a grid floor wired to a shock generator surrounded by an acoustic chamber (Ugo Basile), and controlled by the EthoVision software (Noldus). After head-fixed habituation, mice were placed in the conditioning box and CSs were presented at random times. For the CS+ a pure tone (7.5 kHz) was presented for 20 sec, followed by a 2 sec foot shock (0.6 mA, repeated 3-5 times). For the CS− a broadband noise was presented for 20 sec but was never associated with a foot shock (10-12 times). The inter-trial interval was random selected between 1-2 min.

To test FC recall, mice were placed in a different context (a cylinder-shaped cage with stripes on the walls and a smooth floor), and then presented with the CSs. Each CS was presented for 5 sec and the inter-trial interval was 25 sec (repeated 5 times). Conditioned freezing was measured during the 5 sec presentation of the CS (and averaged over trials). Freezing was automatically measured throughout the testing trial by the EthoVision tracking software. For head-fixation recall, the same CSs were presented (5 sec long, 25 sec inter-trial-interval, repeated 18-24 times) and freezing was measured by pupil diameter.

### Immunostaining

For immunostaining of LC neurons and NE axons, 50-μm thick coronal sections were collected sequentially into phosphate buffered saline (PBS). Then, sections were washed in PBS and blocked for 2–3 h at room temperature in 10% normal donkey serum (NDS) in PBS with 0.3% Triton-X100 (PBST). For immunostaining of LC neurons, primary antibodies for tyrosine hydroxylase were used (rabbit anti-tyrosine hydroxylase, Millipore, AB152, 1:2,000). For immunostaining of NE axons in the cortex, primary antibodies for noradrenaline transporter were used (mouse anti-noradrenaline transporter was used, PhosphoSolutions, 1447-NET, 1:10000; rabbit anti-GFP, novusbio NB600-308, 1:1,000). The primary antibodies were diluted in 5% NDS in PBST and incubated overnight at room temperature with mild shaking. After 3×10 min washes in PBST, secondary antibodies (donkey anti-rabbit, Alexa-488, 1:500 and donkey anti-mouse, Alexa-647, 1:500, Jackson ImmunoResearch) were applied for 1– 2 h at room temperature with mild shaking, followed by 3×10 min washes in PBST. The sections were mounted and imaged with a confocal microscope (FV-10i - Olympus). Images were processed using ImageJ software.

### Head-fixed sensory stimuli

Whiskers were stimulated by air puffs for a duration of 200-500 ms. Air puffs were delivered from a compressed air tank to a tube ending with a pipette tip facing the mouse’s whiskers. For auditory stimulation a broadband noise was presented with speakers for 100 ms. Visual stimulation was delivered by presenting light flashes with a green LED for 200-500 ms.

### Pupil

Recording of pupil diameter was performed using a camera (ELP-USBFHD01M-L36-120FPS) at 20 Hz. A moderate level of ambient illumination was maintained by 410-420 nm high power LEDs (743-IN-K2PUV-U70, Inolux, Mouser Electronics). Pupil segmentation was performed with SimplePupil - a LabVIEW-based pupil tracker for neuroscience applications developed in Ilan Lampl’s Lab.

The evoked change in pupil diameter, ΔD/D_0_ (percent) was calculated as ((D - D_0_) / D_0_) * 100, where D_0_ is the mean activity during the 2 sec preceding the stimulus. The mean evoked response was calculated as the mean ΔD/D_0_ during the 10 sec following stimulus onset.

### Data Analysis

#### Ca^2+^ Imaging

All analyses were performed using ImageJ software (Schneider et al., 2012) and custom-written codes in Python. Movements were corrected using the moco plugin (Dubbs et al., 2016). ROIs were manually selected, and fluorescence traces were low pass-filtered at 3 Hz. The evoked fluorescence change, ΔF/F_0_ (percent) was calculated as ((F - F_0_) / F_0_) * 100, where F_0_ is the mean activity during the 1 sec preceding the stimulus. In spontaneous fluorescence traces, F_0_ was defined as the tenth percentile of F. The mean evoked response was calculated as the mean ΔF/F_0_ during the 2 sec following stimulus onset. To avoid the possibility of visual stimulation affecting the imaging fluorescence, in figure 2 the mean evoked response was calculated as the mean ΔF/F_0_ only starting 0.5 sec following stimulus onset for all conditions. In figure 3I the mean evoked response was calculated as the mean ΔF/F_0_ during the entire auditory stimulation (5 sec). All stimulation occurring immediately before or during running periods were excluded from the analysis.

We defined a response to a stimulation if the mean evoked response was larger than 8% fluorescence (changing the threshold yielded similar results). The response probability was defined as the mean of responses over all trials. Bootstrap was done by shuffling the times of stimulations.

In figure 4E-F, dendrites that responded in more than 40% of the trails were selected for analysis. In figure 4E, ‘weak axonal response’ was defined as the mean evoked response in trials were the axonal response was in the first quartile of responses, while the ‘strong axonal response’ was defined as the mean evoked response in trials were the axonal response was in the fourth quartile of responses. Each axon was discretized separately. In figure 4F axonal responses were discretized based on the deciles.

### Statistical Analysis

Statistical significance was assessed using an unpaired Student’s two-sided t test. In figure 3B,F,I, and figure 4E a paired t test was used. In figure 2C-D, p values were computed using analysis of variance (ANOVA) with Tukey post-hoc test. Significance level were marked as *p < 0.05, **p < 0.01, and ***p < 0.001. All data are reported as mean ± S.E.M unless otherwise specified.

